# Visual confidence accurately tracks increasing internal noise with eccentricity in peripheral vision

**DOI:** 10.64898/2026.01.28.702447

**Authors:** Luhe Li, Michael S. Landy

## Abstract

Sensory representations are inherently noisy, and monitoring this noise is essential for effective decision-making. This metacognitive ability of evaluating the quality of one’s perceptual decision is referred to as perceptual confidence. However, whether perceptual confidence accurately tracks internal noise remains unresolved. Peripheral vision provides a natural testing ground for this question, yet previous studies report mixed results complicated by different definitions and measurements of confidence. Here, we used a normative Bayesian framework with incentivized confidence measurements to address these discrepancies. We tested the Bayesian-confidence hypothesis that confidence is derived from the posterior probability distribution of the feature being judged, given noisy sensory measurements. We tested two perceptual tasks while varying stimulus eccentricity: spatial localization and orientation estimation. We measured confidence by post-decision wagering, by which participants set a symmetrical range around the perceptual estimates. Participants earned higher reward for narrower confidence ranges but received zero reward if the range did not enclose the target. We estimated sensory noise from the perceptual responses to predict confidence, assuming that sensory noise linearly increases with eccentricity. We then compared a normative Bayesian model with three alternative models that challenged different assumptions. Across both tasks, the Bayesian ideal-observer model best predicted confidence. These results suggest that humans can accurately monitor the increased internal noise in peripheral vision and use this information to make optimal confidence judgments.

## 1 Introduction

Sensory representations are inherently noisy [1]. Consequently, effective decision-making based on these percepts requires monitoring this noise. When a driver checks a rain-streaked side mirror or a pedestrian navigates a foggy crossing, the decision-maker must infer not only what and where things are in the world, but also how reliable those percepts are for guiding subsequent action. This metacognitive ability of evaluating the quality of one’s perceptual decision is referred to as perceptual confidence [2]. Confidence plays a crucial role in the planning of subsequent actions after a decision [3, 4], learning [5], and cooperation in group decision making [6].

The two relevant concepts, noise and confidence, are not monolithic. A distinction must be made between external noise, which arises from variability in the environment, and internal noise, which is inherent to the perceptual system [7, 8]. It is well established that humans account for internal noise in cross-modal integration [9–11], recalibration [12–14], and sensorimotor learning [15], often more effectively than for external noise [8]. Whether this ability extends to metacognition remains unknown—specifically, whether perceptual confidence accurately tracks internal noise. Resolving this question is essential for a mechanistic understanding of how an observer navigates perception in noise through metacognitive evaluation.

Prominent process models of confidence posit that confidence should track internal noise. For example, the extended Signal Detection Theory (SDT) computes confidence from the same sensory signal and noise that govern the first-order perceptual decision [16–19]. Similarly, the Bayesian-confidence hypothesis [20] proposes that confidence is derived from the posterior probability distribution of the feature being judged, given noisy sensory measurements [21–25]. Some empirical results have reported behaviors consistent with this hypothesis [25–27].

Meanwhile, other results suggest that confidence may incorporate sensory noise in a probabilistic yet non-Bayesian manner, such as via stimulus-response mapping [20, 28–31]. Additional evidence shows that confidence can be driven by heuristic cues rather than noise, such as the magnitude of the sensory measurement [32], reaction time [33], task-difficulty variables [34], and post-decision processing [35, 36].

These accounts make different and testable predictions about how confidence should change as internal noise varies in a known way. Peripheral vision is ideal for this test because internal noise increases systematically with eccentricity and has been extensively quantified across domains, such as acuity [37–39], contrast sensitivity [40–46], color sensitivity [47–49], and crowding [50–54]. Yet studies of confidence in peripheral vision have reached divergent conclusions, largely because confidence has been defined and measured in different ways. Even when objective performance is carefully equated across eccentricities, prior work has reported overconfidence (“subjective inflation”; [55–58]), underconfidence [59], and accurate confidence in peripheral judgments [60].

To address these discrepancies, we combined the normative Bayesian framework with an incentivized continuous post-decision wagering task. Participants performed spatial-localization or orientation-estimation tasks across varying eccentricities and reported confidence by setting a range around their perceptual estimates. A smaller confidence range was associated with a higher reward, but missing the target resulted in zero reward.

This approach offers three key advantages over previous categorical confidence self-reports (e.g., Likert scale or binary low/high confidence decisions). First, it uses the well-accepted definition of confidence, the subjective probability that a perceptual decision is accurate [21, 24, 26]. Second, measuring confidence by post-decision wagering in the same units as the first-order task offers interpretable and incentivized confidence responses [61–63]. Third, by comparing a normative Bayesian model against alternative models, we do not necessarily assume optimality. Rather, we use the ideal observer as a benchmark to pinpoint the specific sources of potential suboptimality in confidence [23, 64, 65].

To preview our behavioral results, we confirmed that the variability of localization and orientation-estimation responses increased with eccentricity, whereas confidence decreased. We estimated sensory noise from the perceptual responses to predict confidence. We compared a normative Bayesian model with three alternative models that challenged the model assumptions. Formal model comparison revealed that the Bayesian ideal-observer model best predicted confidence data across both first-order tasks. These results suggested that despite the noisy nature of the perceptual system, observers accurately monitor their increasing internal noise with eccentricity to form optimal confidence judgments that maximize expected gain.

## 2 Results

### 2.1 Experiment 1

#### 2.2.1 Behavioral results

We measured confidence in a spatial-localization task using post-decision wagering (Fig. 1A). In each trial, a Gaussian blob was presented for 33 ms at a random position on the invisible horizontal axis (*±*28 degree visual angle, dva, uniformly sampled; Fig. 1A). Participants were instructed to fixate crosshairs in the center of the display until the target disappeared, after which they could move their eyes freely. We used an eye-tracker (EyeLink 1000, SR Research Ltd., Ottawa, Canada; sampling rate: 1000 Hz) to monitor participants’ gaze position, and skipped and repeated trials if the gaze shifted away from the crosshairs center more than 2 dva during stimulus presentation. In the response period, participants adjusted the horizontal position of a cursor to the perceived center location of the target using a mouse roller-ball and clicked to confirm the location estimate. Next, they turned a dial that symmetrically widened a confidence range around their location response while linearly reducing the potential reward (0.01–1 points per trial, reducing to 0.01 when the range was wider than one-fourth of the full screen width). If the confidence range failed to enclose the target center, the actual reward was zero (upper panel, Fig. 1B); if it enclosed the target center, participants earned the potential reward (lower panel, Fig. 1B). No trial-wise feedback was given so that wagers reflected participants’ estimates of internal noise rather than learned task-specific strategies [29].

**Figure 1.**
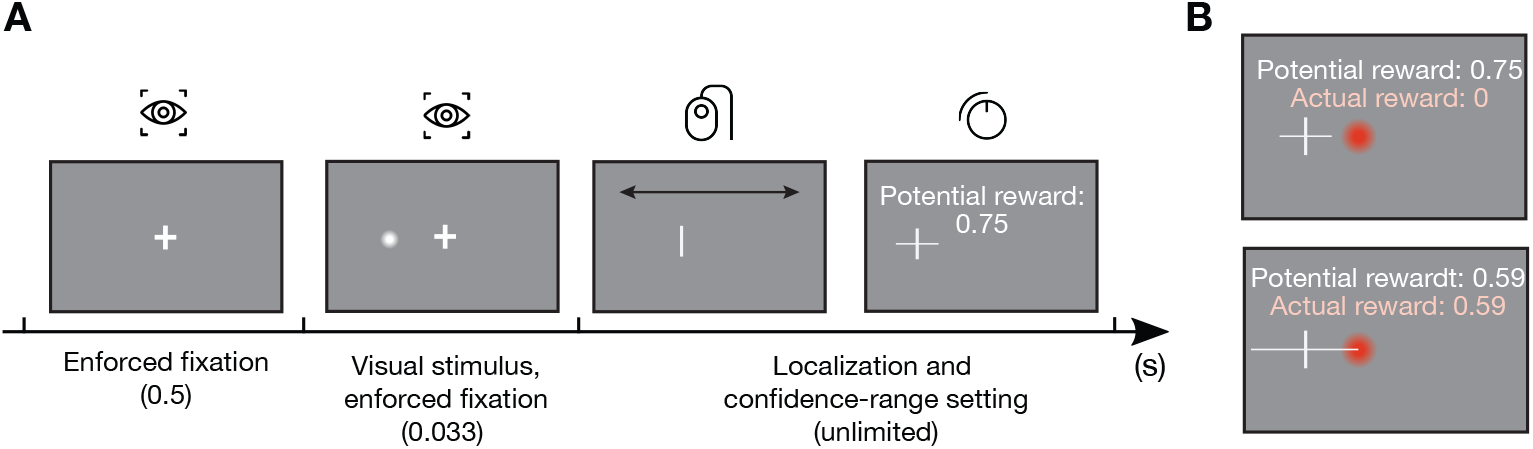
Experiment 1, spatial localization and confidence task. (A) Task timing. In each trial, participants fixated crosshairs at the center of the display. Then, the stimulus (a white Gaussian blob) appeared at a uniformly sampled random location on the invisible horizontal axis. After the stimulus disappeared, participants adjusted the cursor’s horizontal location with a mouse rollerball to reproduce the center location of the target, and set a confidence range symmetrically placed around the location estimate with a dial to earn a reward (0.01–1 points per trial). Increasing the confidence range linearly decreased the potential reward. (B) Example of the reward in the confidence range-setting task. Only elements in white were visible to participants. Upper panel: The actual reward is zero because the confidence range is too narrow to include the center of the target (blob shown in red for illustration). Lower panel: The actual reward is the same as the potential reward when the confidence range encloses the target center.

An example participant’s raw data show two key features of the localization responses (Fig. 2A). First, localization variability increased with eccentricity. We binned trials by eccentricity and calculated the standard deviation (SD) of localization responses in each bin, and found a monotonic increase as a function of eccentricity (Fig. 2B). Second, the localization responses were systematically pulled towards the center, suggesting a central spatial bias. A linear fit of reproduced versus target locations yielded slopes less than one at the group level (mean slope = 0.96 ± 0.01; one-sample *t*(14) ≈ –3.51, *p* = 0.002), with one participant showing a bias towards the periphery.

**Figure 2.**
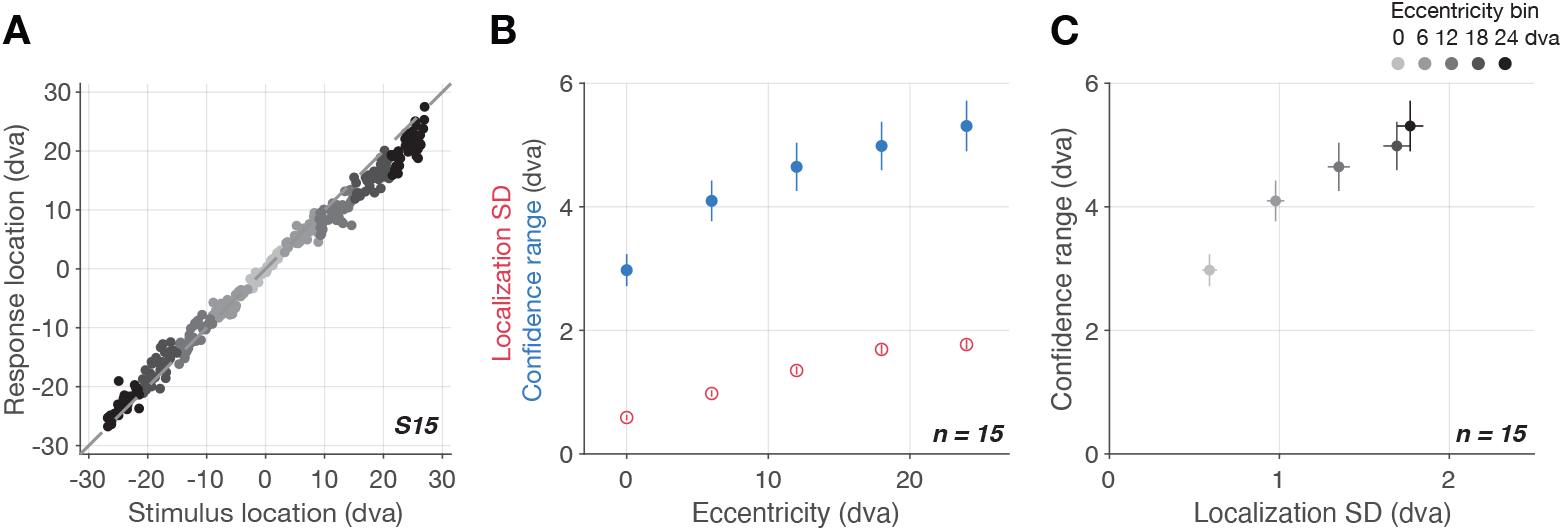
Behavioral results. (A) Localization responses of an example participant. Darker colors represent larger eccentricity. Localization variability increased with eccentricity. (B) Increase of localization standard deviation (SD) and confidence range with eccentricity. A larger confidence range indicates lower confidence. Error bars: *±*1 SEM across participants. (C) Increase of confidence range as a function of localization SD. Vertical and horizontal error bars: *±*1 SEM across participants.

More importantly, the confidence range tracked the localization variability well. The larger the eccentricity, the larger the confidence range, i.e., less confidence (Fig. 2B, C). Although this behavioral pattern was clear, only comparison of generative models can reveal the sources and computations of confidence. We therefore evaluated several candidate confidence models.

#### 2.1.2 Modeling results

##### Localization models

All confidence models build on the same localization model. This model assumes that sensory noise increases with eccentricity, resulting in an asymmetric likelihood function. For each observer, we compared four localization models that differed in how the central spatial bias was implemented and what estimator was used to read out the posterior (SI Appendix 1: Comparing spatial-localization models). We then determined internal noise parameters using the best-fitting model for each observer.

##### Confidence models

In this section, we start by describing the Bayesian ideal-observer model of confidence, followed by reduced models that relax its key assumptions. In the second-order confidence task, the observer sets a symmetric confidence interval by adjusting the confidence range around the first-order estimate (i.e., the reproduced location). Widening the confidence range increases the probability of accurately capturing the target center, reflected by the increased cumulative distribution function (CDF) of the posterior belief of stimulus locations (green line, Fig. 3A). An increase in the confidence range also linearly decreases the reward until it reaches the lowest possible point (purple line, Fig. 3A). An Bayesian ideal-observer multiplies the posterior distribution over location from the localization task with the subjective-gain function to obtain the expected gain function (black line, Fig. 3A), and selects the optimal confidence range (black star, Fig. 3A) by maximizing expected gain. Finally, this optimal confidence range is corrupted by confidence noise [66, 67], producing the confidence-range response.

**Figure 3.**
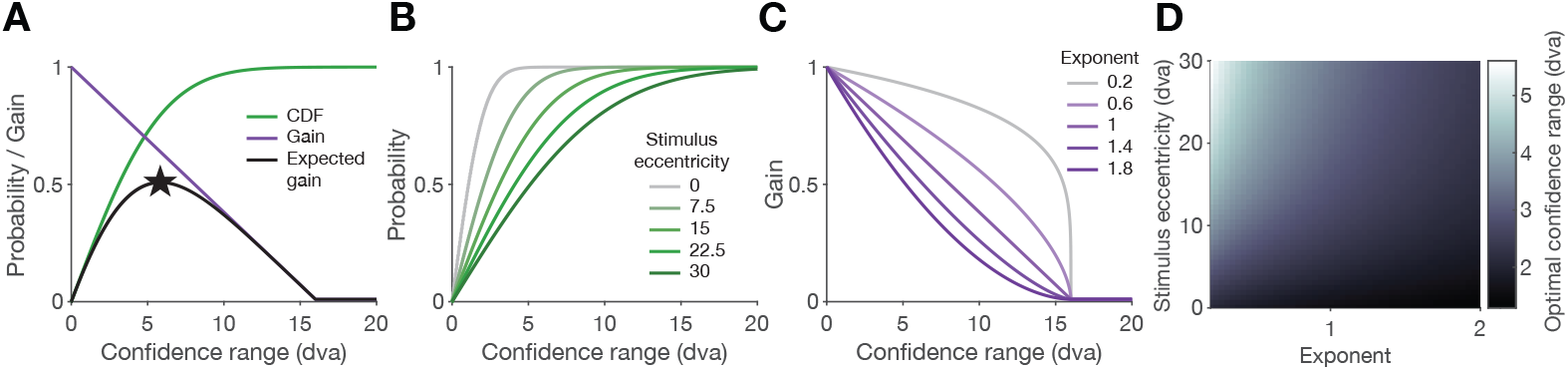
The ideal-observer confidence model. (A) Effects of the confidence range. A larger confidence range increases the probability of enclosing the target (green: cumulative distribution function, CDF, of the posterior belief of stimulus locations), but decreases the potential reward (purple: subjective-gain function). An Bayesian ideal-observer computes their product to obtain the expected-gain function (black) to select the optimal confidence range (black star) that maximizes expected gain. Parameters for example figures: eccentricity = 15 dva, exponent = 1, noise = 0.1*·* eccentricity + 0.8. (B) Effect of the stimulus eccentricity. A larger stimulus eccentricity is associated with larger sensory noise, which reduces the slope of the posterior CDF. (C) Effect of the exponent of the subjective-gain function. Increasing the exponent value changes the shape of the subjective-gain function from concave to convex, which decreases the optimal confidence range and implies more risk-seeking decisions. (D) Effects of the stimulus eccentricity and the exponent. The optimal confidence varies nonlinearly as a joint function of stimulus eccentricity and the exponent of the subjective-gain function.

The optimal confidence range depends on two parameters. Stimulus eccentricity influences the confidence model by modulating the posterior distribution that governs the localization model. For example, a stimulus presented in the periphery is associated with greater internal noise, resulting in a shallower CDF of the posterior (darker lines, Fig. 3B). Consequently, a larger confidence range is required to achieve the same probability of enclosing the target.

Another parameter that influences the resulting confidence range is the exponent of the subjective-gain function, which controls its convexity. The exponent value reflects the risk tolerance of the observer: a value larger than one produces a convex subjective-gain function, leading to a narrower and more risky choice of optimal confidence range (darker lines, Fig. 3C). By contrast, a value less than one represents a risk-averse observer who tends to choose a wider range (lighter lines, Fig. 3C). These two parameters have intuitive implications. For example, a risk-averse observer (i.e., one with a smaller exponent) is more likely to adopt a larger confidence range when the stimulus appears in the far periphery (i.e., at larger eccentricities). More critically, a heatmap of the optimal confidence range as a joint function of stimulus eccentricity and the exponent reveals that the relationship is not a linear combination of these two factors (Fig. 3D). The increase of confidence range with eccentricity is more pronounced when the exponent is small, whereas it becomes relatively flat when the exponent is large.

We tested three alternative models that examine the assumptions of the ideal-observer model (Table 1). First, we evaluated whether the posterior distribution is essential to derive the confidence range. The observer can rely on low-level sensory cues as heuristic strategies instead of performing full Bayesian inference [20, 31, 32]. We tested a heuristic model, the scaled-measurement model, where the confidence range is based on the magnitude of the location measurement. For example, an observer might use the size of the post-stimulus saccade via an efference copy as a heuristic cue for confidence.

**Table 1.**
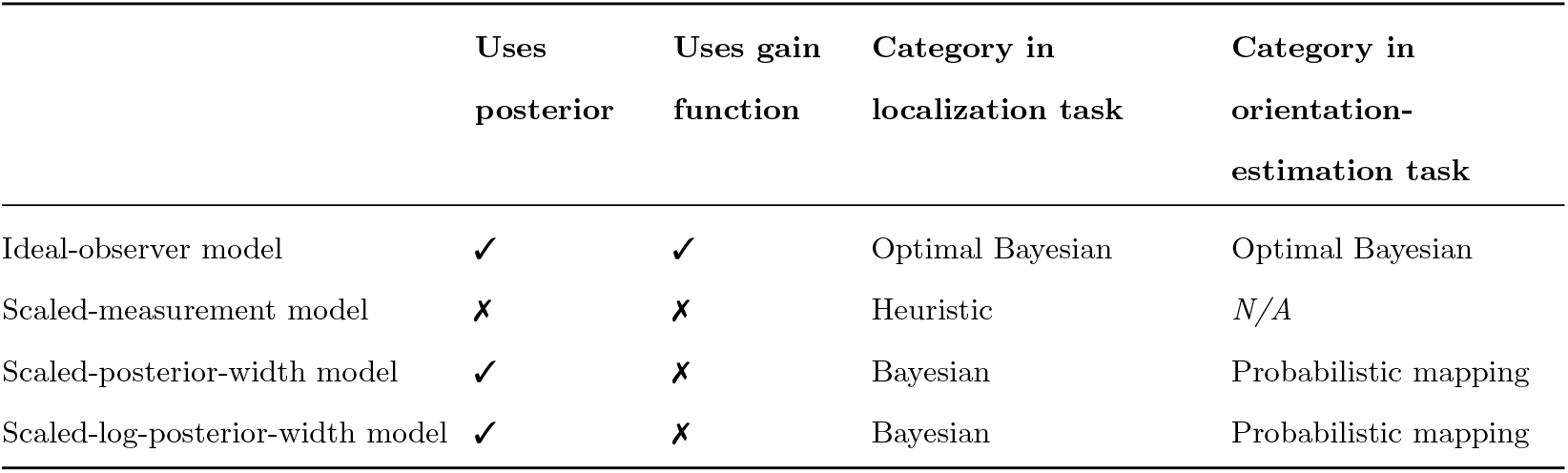
Assumptions implemented in each confidence model. “Use posterior” indicates that the model derives confidence from the posterior distribution over the potential stimulus feature. “Use gain function” indicates that the model maximizes expected gain. “Category” indicates the nature of the confidence model given the specific first-order task. The last two models reduces to non-Bayesian probabilistic-mapping models in the orientation-estimation task because of a flat prior and a symmetric likelihood.

|  | Uses<br>posterior | Uses gain<br>function | Category in<br>localization task | Category in<br>orientation-estimation task |
| --- | --- | --- | --- | --- |
| Ideal-observer model | ✓ | ✓ | Optimal Bayesian | Optimal Bayesian |
| Scaled-measurement model | ✗ | ✗ | Heuristic | $N/A$ |
| Scaled-posterior-width model | ✓ | ✗ | Bayesian | Probabilistic mapping |
| Scaled-log-posterior-width model | ✓ | ✗ | Bayesian | Probabilistic mapping |

Second, we evaluated the assumption that the observer uses the subjective-gain function to obtain the confidence range. We tested two alternative Bayesian models that assume the confidence range is a monotonic function of the posterior width, which is a natural quantification of confidence as a continuous variable [22]. The scaled-posterior-width model scales the posterior width by a constant. A Fechnerian model, the scaled-log-posterior-width model scales the log of the posterior width by a constant. Such a Fechnerian model accounted for discrete confidence reports well in a word recognition-memory task [68].

##### Model predictions

We fit the localization model to each participant’s localization data by maximizing likelihood first, and then fixed the parameters that determine sensory noise for each individual before fitting the confidence models to that individual’s confidence data.

The ideal-observer model captures the increase of confidence range with eccentricity (left-most blue panel, Fig. 4A). Both the heuristic scaled-measurement model and the scaled-posterior-width model exhibit a systematic bias: they underpredict confidence ranges for small eccentricities and overpredict them for large eccentricities. This underprediction is more prominent in the scaled-measurement model, because confidence, as a scaled function of the location measurement magnitude, approaches zero when the stimulus is near the center. Finally, the scaled-log-posterior-width model yields similar model predictions to the ideal-observer model, which also captures the data well. Individual-level model predictions are reported in SI Appendix 2.

**Figure 4.**
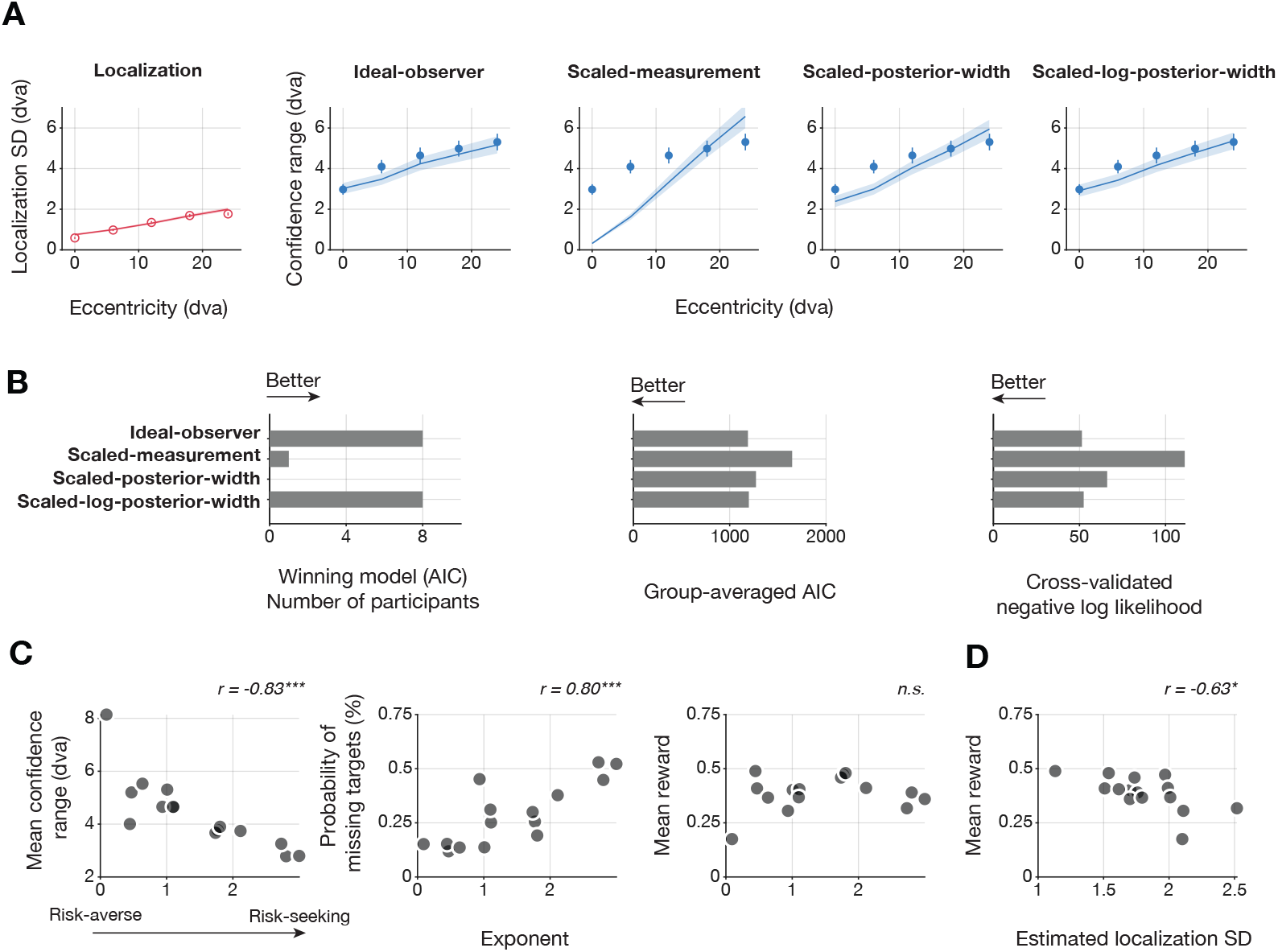
Model predictions and model comparison of Experiment 1. (A) Group-averaged data (dots, *n* = 15) and model prediction (lines and shades) of the localization model and each confidence model. The localization-model prediction (red) is the same across confidence models. The ideal-observer model (left-most blue panel) and the scaled-log-posterior-width model (right-most blue panel) best capture the confidence data. Error bar and shades: ±1 SEM across observers. (B) Quantitative metrics of confidence models. The ideal-observer model and the scaled-log-posterior-width model capture the largest number of participants based on the Akaike information criterion (AIC), have the least group-averaged AIC, and the least five-fold cross-validated negative log likelihood. (C) Correlation of the behavioral measurements with the estimate of the exponent (the risk-tolerance parameter) from the ideal-observer model. Each dot is an observer. Left panel: Larger exponent values (higher risk tolerance) are associated with a smaller confidence range across observers. Middle panel: Larger exponent values are associated with a higher probability of missing the target. Right panel: The exponent does not correlate with the trial-averaged reward. ∗ ∗ ∗ : *p <* 0.001. (D) Correlation between sensory noise and performance. Higher localization SD estimated using the localization model correlates with lower trial-averaged reward. ∗ : *p <* 0.05.

##### Model comparison

Next, we compared the confidence models quantitatively using two metrics. First, based on Akaike information criterion (AIC, [69]), the majority of participants were best fit by either the ideal-observer model or the scaled log posterior width model (left panel, Fig. 4B). The group-averaged AIC also supported that these two models outperformed the others (middle panel, Fig. 4B). Second, we split the data randomly into five folds and computed cross-validated negative log likelihood averaged across folds (right panel, Fig. 4B). The same two models best predicted the data. Therefore, consistent with model predictions, the ideal-observer model and the scaled-log-posterior-width model achieved the best and comparable goodness-of-fit.

##### Parameter estimates

Although the two winning models were statistically indistinguishable, they differ in interpretability. The scaled log-posterior width model lacks a clear mechanistic justification for its logarithmic transform or scale factor. By contrast, the ideal-observer model defines a process-level free parameter: the exponent of the gain function. An exponent greater than one implies risk-seeking decision-making, which is consistent with the parameter estimates across observers. Larger exponent values (higher risk tolerance) are associated with a narrower trial-averaged confidence range (i.e., higher confidence; left panel, Fig. 4C; *r*(13) = −0.83, *p <*.001) and a higher probability of missing the target (middle panel, Fig. 4C; *r*(13) = 0.80, *p <*.001). While participants have different risk tolerance, their trial-averaged reward does not significantly correlate with the exponent value (right panel, Fig. 4C, *r*(13) = −0.18, *p* = 0.52). Instead, the reward is correlated with the estimates of localization SD, which depend on the strength of the linear transformation between the eccentricity and sensory noise in the localization model (Fig. 4D; *r*(13) = −0.63, *p* = 0.012). Observers with more sensory noise tend to receive fewer rewards. Individual-level parameter estimates of the localization model and winning confidence models are reported in SI Appendix 3.

### 2.2 Experiment 2

#### 2.2.1 Behavioral results

To test the generalizability of the confidence model, we used an orientation-estimation task as the first-order task in Experiment 2. The orientation-discrimination task has been widely used in previous studies of confidence in peripheral vision (e.g., [23, 27, 57, 59, 70, 71]).

In each trial, participants fixated on a central dot while a Gabor patch with a random orientation (0–*π*, uniformly sampled, 0 is horizontal) appeared at a random position along an invisible horizontal axis (*±*28 dva, uniformly sampled; Fig. 5A). Gaze position was monitored by the eye-tracker to ensure fixation until stimulus offset. Participants then turned a dial to rotate a response bar presented at the same location as the stimulus with a random initial orientation to match the perceived orientation. The confidence task was identical to that in Experiment 1, except that participants turned the dial to adjust the width of a confidence wedge symmetrically centered on their orientation estimate.

**Figure 5.**
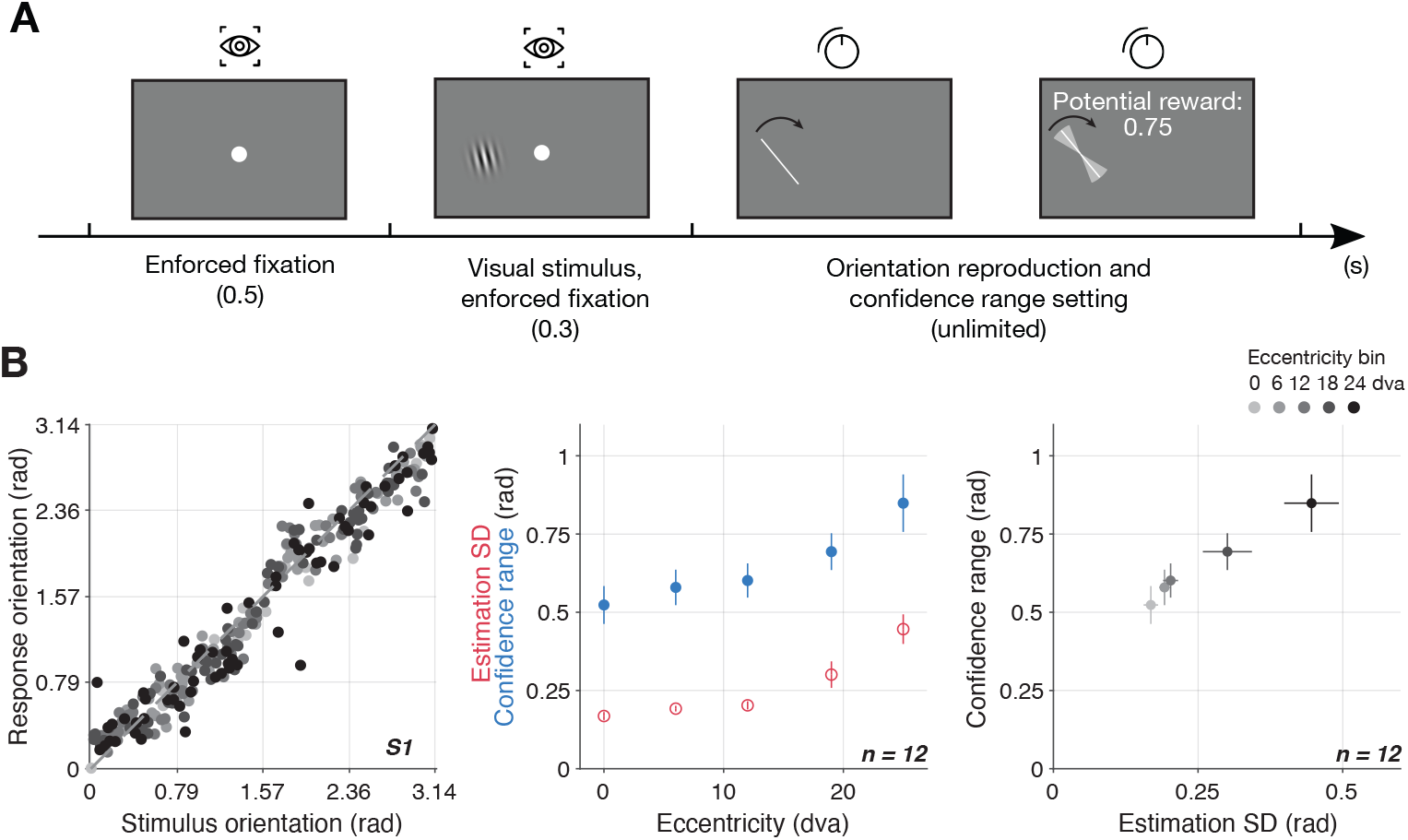
Experiment 2, Orientation-estimation and confidence task. (A) Task timing. In each trial, participants fixated a circle in the center of the display. Then, a stimulus with a random orientation appeared at an independently-sampled random location on the invisible horizontal axis. After the stimulus disappeared, they used a dial to rotate a response bar at the stimulus location to reproduce the target orientation and then used that same dial to set a confidence wedge symmetrically placed around the orientation estimate to earn a reward (0.01–1 points per trial). (B) Behavioral results. Left panel: Orientation-estimation responses of an example participant. Darker colors represent larger eccentricity. Middle panel: Increase of orientation-estimation SD and confidence range with eccentricity. A larger confidence range indicates a lower confidence level. Error bars: ±1 SEM across participants. Right panel: Increase of confidence range as a function of orientation-estimation SD. Vertical and horizontal error bars: *±*1 SEM across participants.

As expected, orientation estimates became noisier with increasing eccentricity (left panel, Fig. 5B). When we binned trials by eccentricity, we confirmed that the SD of orientation estimates increased, while confidence decreased, in the periphery (middle and right panels, Fig. 5B).

#### 2.2.2 Modeling results

##### Orientation-estimation model

All confidence models build on the same orientation-estimation model, which rests on only two assumptions. First, sensory noise linearly increases with eccentricity. An alternative model in which sensory noise is an exponential function of the eccentricity yields the same results in terms of preferred confidence model (SI Appendix 4). Second, the probability density function (PDF) of the orientation measurement follows a von Mises distribution, whose concentration parameter is determined by sensory noise. Unlike the localization model, the likelihood function is symmetric because all possible orientations within a trial share the same sensory noise determined solely by the stimulus eccentricity in that trial. The orientation-estimation model fit the estimation responses well (Fig. 6A), allowing us to estimate sensory noise for all the confidence models.

**Figure 6.**
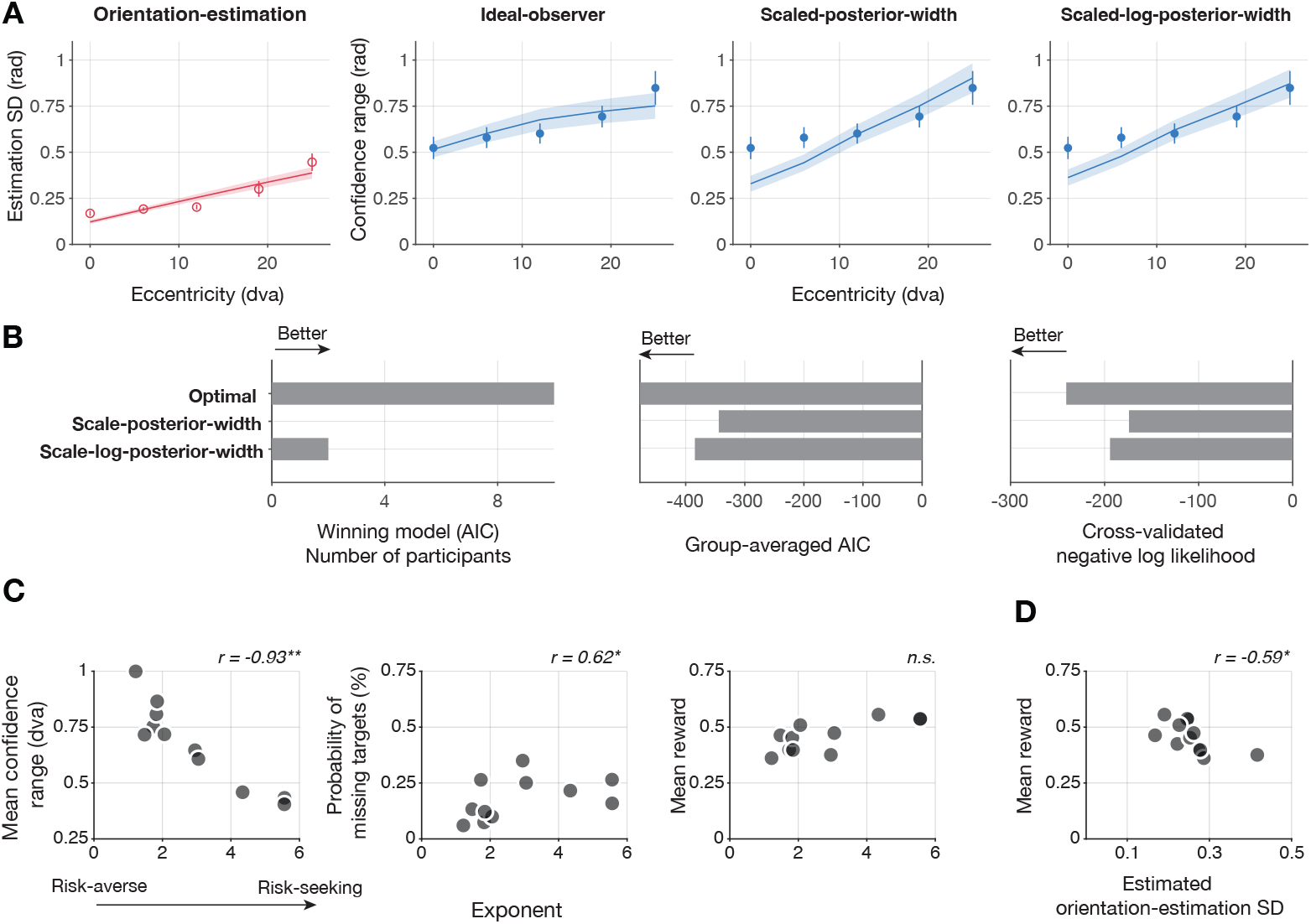
Model predictions and model comparison for Experiment 2. (A) Group-averaged data (dots, *n* = 12) and model prediction (lines and shades). The orientation-estimation model prediction (red) is fixed across all panels. The ideal-observer model (left-most blue panel) best captures the confidence data. Error bar and shades: ±1 SEM across observers. (B) Quantitative metrics of confidence models. The ideal-observer model captures the majority of participants based on AIC, has the least group-averaged AIC, and the least five-fold cross-validated negative log likelihood. (C) Correlation of the behavioral measurements with the estimate of the exponent (the risk-tolerance parameter) from the ideal-observer model. Each dot represents one observer. Left: Higher risk tolerance is associated with a smaller confidence range. Middle: Higher risk tolerance is associated with a higher probability of missing the target. Right: No significant correlation exists between the exponent and trial-averaged reward. ∗: *p <* 0.05, ∗∗: *p <* 0.01. (D) Correlation between sensory noise and performance. Higher orientation-estimation SD estimated by the orientation-estimation model correlates with lower trial-averaged reward.

##### Confidence models

The Bayesian ideal-observer model of confidence is identical to that in Experiment 1 (Table 1). The alternative confidence models are a subset of those tested in Experiment 1 because we did not include the scaled-measurement model. Using task-irrelevant location information would require embedding the full localization model in the orientation-estimation model, adding three free parameters and increasing overfitting risk. This model will also underpredict confidence in the center because the location measurement approaches zero at the fovea, as shown in the model predictions of Experiment 1 (Fig. 4A). We therefore tested only the remaining three confidence models for orientation estimation. Furthermore, the scaled-posterior-width model and the scaled-log-posterior-width model reduce to the non-Bayesian probabilistic mapping model when the likelihood is symmetric and the prior is uniform. In other words, they are equivalent to models that directly transform the estimates of sensory noise.

##### Model predictions and model comparison

The ideal-observer model better captures the confidence responses, whereas the other two models consistently underpredict the confidence at low eccentricity. Unlike Experiment 1, the scaled-log-posterior-width model deviates from the ideal-observer model predictions and fails to account for confidence in this task for most of the observers (SI Appendix 5: individual-level model predictions). Quantitatively, the ideal-observer model captures the largest number of participants by AIC, yields the lowest group-averaged AIC, and achieves the lowest five-fold cross-validated negative log likelihood. Taken together, the same ideal-observer model captures confidence well across two first-order tasks.

##### Parameter estimates

The risk-tolerance parameter also plays similar roles in both experiments. The exponent of the subjective-gain function is negatively correlated with the trial-averaged confidence range (*r*(10) = −0.93, *p <*.001) and positively correlated with the probability of missing the target (*r*(10) = 0.62, *p <*.05). These results suggest that more risk-seeking participants are more likely to set a smaller confidence range and to miss the target. Despite the different risk tolerance, the trial-averaged reward does not differ significantly across participants (*r*(10) = 0.54, *p* = 0.07), but is correlated with orientation-estimation variability (*r*(10) = −0.59, *p <*.05). Individual-level parameter estimates of the ideal-observer model are reported in SI Appendix 6.

## 3 Discussion

We examined whether perceptual confidence accurately tracks increasing internal noise with eccentricity in peripheral vision. We quantified how variability in spatial localization and orientation estimation increased with eccentricity, and used these estimates of internal noise to predict confidence measured by a continuous post-decision wagering paradigm. Model comparison revealed that an ideal-Bayesian observer, which uses the full posterior distribution based on accurate knowledge of eccentricity-dependent noise to optimize expected gain, provided the best account of confidence in both tasks. The reduced Bayesian model without a subjective-gain function succeeded in only one task, and the heuristic model never provided the best fit. These results support that confidence accurately tracks internal noise governing perceptual decisions and is consistent with predictions of optimal Bayesian inference.

### 3.1 Bayesian vs. non-Bayesian confidence models

We found that confidence in peripheral vision is best explained by an Bayesian ideal-observer model. In the spatial-localization task, both the ideal-observer and a model that scales the log of posterior width utilized the full posterior distribution and fit the data well. However, in the orientation-estimation task, the scaled-log-posterior-width model simplified to a direct transformation of a scalar estimate of sensory noise and failed. This dissociation reveals that a summary statistic of sensory noise is insufficient; accurate confidence judgments require integrating over the posterior. Notably, the Bayesian ideal-observer model predicted confidence in both tasks despite their distinct likelihood structures, suggesting a common mechanism for uncertainty-based confidence judgments.

Our results add to the evidence supporting the Bayesian-confidence hypothesis [20, 22, 25, 70], suggesting that observers can make optimal confidence judgments when sufficiently incentivized. However, this does not imply that confidence is universally Bayesian. Rather, it likely reflects a resource-rational trade-off between computational limits and potential reward [23, 72]. For example, high computational costs of three distinct options may prompt a switch to suboptimal strategy [28]. Consequently, we advocate using incentivized tasks and comparing ideal-observer models against reduced and heuristic alternatives to pinpoint when and how confidence deviates from optimality.

### 3.2 Confidence tracks internal noise

Our model aligns with a broad class of SDT models in which confidence and the perceptual decision arise from the same sensory evidence [16, 17, 19, 73, 74]. These models differ in terms of how the sensory evidence is subsequently corrupted by metacognitive noise. While we assumed log-normal confidence noise, other noise formulations (reviewed in [75]) could be incorporated within the same framework.

More broadly, our results are consistent with recent proposals conceptualizing confidence as decision consistency or reliability given the sensory evidence [71, 76]. This view, formalized in the CASANDRE model, posits that observers must estimate decision reliability based on sensory noise, because they lack trial-by-trial feedback of decision accuracy [77]. Our Bayesian formulation naturally incorporates trial-by-trial estimates of sensory noise into the posterior distribution. Thus, the Bayesian definition reconciles confidence as both perceived accuracy (a belief instead of ground-truth) and decision consistency.

Our experimental results corroborated that confidence is calibrated to internal noise. While studies that manipulated internal noise have reached mixed conclusions regarding this calibration (e.g., [27, 70, 78, 79] vs. [30, 80, 81]), we demonstrate clear internal-noise monitoring when exploiting the natural variation of eccentricity. Importantly, our model does not assume a specific noise profile (i.e., that noise must always increase with eccentricity). For visual attributes where sensitivity retains or increases in the periphery (e.g., temporal-frequency discrimination; [82]), we predict that confidence will lawfully track this trend of internal noise and can be higher in the periphery than in the fovea.

### 3.3 Towards reconciling conflicting accounts of confidence in peripheral vision

Compared with previous work on confidence in peripheral vision, our approach differs mainly in two aspects. First, we measured confidence using a continuous wagering report rather than discrete rating scales that can vary across observers and across time [2]. Because the confidence response is continuous and expressed in the same units as the first-order estimate, it also yields a richer and more interpretable dataset for model fitting. Second, we did not follow the classic approach of equating objective performance (*d*^*′*^) across eccentricity conditions. Although equating *d*^*′*^ facilitates model-free comparisons, it requires stimulus manipulations (e.g., increased contrast, size or spatial frequency) that introduce confounding heuristic cues. By leveraging process models, we avoided these confounds and examined the sources and computations of confidence. Despite methodological differences, our finding of Bayesian optimality provides a mechanistic explanation for recent reports of stable metacognitive efficiency from the fovea to up to 40 dva [60].

Our results do not support the account of overconfidence in the periphery [55–58], which defines confidence as a more liberal criterion. Such overconfidence has been explained by an SDT model that postulates that the observer uses a fixed criterion for detecting targets in high- and low-noise conditions [55]. However, recent work has challenged this view, showing that a flexible criterion that scales with sensory noise can also account for these empirical results [83]. While our definition of confidence differs from the detection-/discrimination-criterion approach, our modeling results align with the latter view that observers maintain an accurate estimate of their own sensory noise to guide criterion placement.

Regarding reports of underconfidence, Toscani, Mamassian, and Valsecchi [59] used a confidence forced-choice paradigm to separate metacognitive bias from metacognitive sensitivity. Their findings are most comparable to ours in terms of metacognitive sensitivity, where they reported a specific deficit when observers compared central and peripheral judgments. However, without a process model, it is unclear whether this impairment reflects a failure to estimate internal noise locally or a difficulty in comparing noise distributions that differ across eccentricities.

## 4 Materials and Methods

### 4.1 Participants

Fifteen participants were recruited through the New York University Department of Psychology’s paid research-participation system (three males; age: 26.3 *±* 9.7) in Experiment 1. A different group of 12 participants was recruited for Experiment 2 (four males, 21.8 *±* 3.0). They all reported normal or corrected-to-normal vision. All participants provided informed written consent before the experiment and received 1.5 credits or $15/hour as compensation. The study was conducted in accordance with the guidelines of the Declaration of Helsinki and approved by the New York University institutional review board.

### 4.2 Apparatus and stimuli

Participants completed the experiments in a dark room. They were seated 30 cm from a CRT monitor (GDM-5402, Sony Electronics Inc., New Jersey, USA; 1,280*×*960 screen resolution, 39.2 cm width, 60 Hz refresh rate) and placed their head on a chin rest with an eye-tracker mounted above (EyeLink 1000, SR Research Ltd., Ottawa, Canada; sampling rate: 1000 Hz). All stimuli were generated and presented using MATLAB R2024b (MathWorks, Natick, MA) and PsychToolbox-3 [84–86].

In Experiment 1, the visual stimulus was a white Gaussian blob (12.69 *cd/m*^2^, SD: 1.5 dva) on a gray background (2.26 *cd/m*^2^). In Experiment 2, the visual stimulus was a Gabor patch (cosine phase, 3 cycle per dva, SD: 0.375 dva) on the same gray background.

### 4.3 Procedure

#### Experiment 1: Localization task

Participants began each trial by fixating central crosshairs for 500 ms while gaze was monitored continuously. If gaze deviated by more than 2 dva from the center at any point before the stimulus disappeared, the trial was canceled and re-queued to be repeated at the end of the session.

A Gaussian blob was then presented for 33 ms at a random horizontal position (uniformly sampled, ±28 dva). Immediately afterward, both the stimulus and the fixation crosshairs disappeared, and a cursor appeared at a random horizontal position. Participants were free to move their eyes from this point onward. Using a roller-ball mouse, they adjusted the cursor’s horizontal position to indicate the perceived target location. The cursor remained visible until they confirmed the position with a mouse click, at which point the cursor was fixed in place.

Next, participants used a dial with their left hand to set a symmetric confidence range centered on the reproduced location. A narrower range increased the potential reward (displayed above the cursor on a scale from 0.01 to 1), but if the true stimulus center location fell outside the range, the actual reward for that trial was zero. When the confidence range equaled or exceeded one-quarter of the display width, the potential reward was reduced to 0.01. Only potential reward values were shown during the trial; actual rewards were not revealed, so participants could not know on a given trial whether the target had been captured. Cumulative actual reward was displayed only during breaks between experimental blocks. There were 270 trials evenly split into 5 blocks. To increase motivation, a leaderboard of all participants’ final scores was shown at the end of each session. Before starting the main experiment, participants completed an 18-trial practice block with trial-by-trial feedback of the actual reward, to familiarize them with the eye-tracker, the response procedure, and payoff rule. A full session lasted approximately 1.5 hours.

#### Experiment 2: Orientation-estimation task

This task followed the same fixation procedure as Experiment 1. A Gabor patch was presented for 300 ms at a random horizontal position (uniformly sampled, *±*28 dva). The Gabor’s orientation was pseudo-randomized on each trial between 0 and *π* (uniformly sampled; 0 is horizontal).

After stimulus presentation, a straight bar appeared at the same location as the stimulus, initialized to a random orientation. Participants were free to move their eyes while using a dial with their right hand to rotate the bar around its center to match the perceived orientation, confirming their response by pressing the dial. They then adjusted the dial further to set a symmetric confidence wedge centered on the reproduced orientation. The payoff rule was similar to the localization task, dropping to 0.01 for a confidence range of 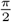 radian. There were 270 trials evenly split into 6 blocks, lasting approximately 1.5 hours.

### 4.4 First-order-task models

#### 4.4.1 Model of spatial localization

We model observers’ behaviors in the first-order task as a Bayesian-inference process for the generative model below. On each trial *i*, the horizontal location of the visual stimulus, *s*_*i*_, is drawn from a uniform distribution, *p*(*s*_*i*_) = 𝒰 (−28, 28) dva.

We assume that the physical stimulus position was linearly remapped into an internal spatial coordinate: 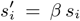, where *β >* 0 captures a spatial bias of visual localization [87]. A model comparison showed that this linear-mapping model outperformed a model that assumed a central spatial prior (SI Appendix 1).

The sensory measurement of location *m*_*i*_ is subject to Gaussian sensory noise,

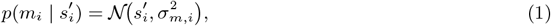

where the sensory noise *σ*_*m,i*_ increases linearly with eccentricity,

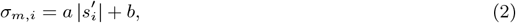

with *a >* 0 and *b >* 0 representing the eccentricity-dependent and baseline components of the noise, respectively. The baseline noise represents the sensory noise at the fovea.

We assume a uniform prior over location, indicating that the observer did not have a prior bias toward particular positions. The likelihood function of the hypothetical stimulus location *s*_*i*_ given a measurement *m*_*i*_ was:

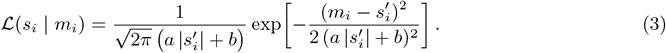

The eccentricity-dependent noise results in an asymmetric likelihood function. This asymmetry arises because, for a given measurement of location, two hypothetical stimuli equidistant from the measured location differ in likelihood: the hypothetical stimulus closer to the fovea is less likely than the one in the periphery because the latter has a wider likelihood function. With a uniform prior, the posterior was proportional to the likelihood and retained its asymmetry.

Next, we consider two estimators that the observer might use to read out the posterior distribution. The maximum-a-posteriori estimator (MAP) corresponds to the mode of the posterior over 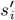. We assume that the response is in the scaled internal space *s*^*′*^ instead of the original space *s*. That is, we assume that the bias in localization of peripheral stimuli is due to an alignment of the internal perceived-location scale (*s*^*′*^) with the allocentric scale used to align gaze with the perceived position of the stimulus when setting the cursor location for the response. The Bayesian-least-squares estimator (BLS) minimizes the mean squared error, corresponding to the mean of the posterior. These two estimates, 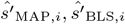, are different because the posterior distribution is asymmetric. We compared models with these two estimators and used the individual-specific best-fitting model for localization.

Given unlimited response time and a motivation to minimize parameters, we model motor noise as a small and fixed constant, *σ*_motor_ = 0.01 dva. The PDF of the observed localization responses *x*_*i*_ is:

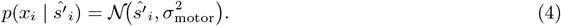

The localization model has three free parameters, Θ_*x*_ = *{β, a, b}*.

#### 4.2.2 Model of orientation estimation

As in the localization task, in each trial *i*, the horizontal location *s*_*i*_ is drawn from the same uniform distribution, *p*(*s*_*i*_) = 𝒰 (−28, 28) dva.

The orientation *θ*_*i*_ is drawn independently from another uniform distribution, *p*(*θ*_*i*_) = 𝒰 (0, *π*) radian, where 0 corresponds to a horizontal grating.

Similarly, we assume the sensory noise (i.e., the circular SD) of orientation measurements is an affine transformation of the eccentricity,

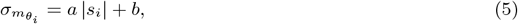

where *a >* 0 and *b >* 0.

The PDF of the orientation measurement, 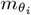, is modeled as a von Mises distribution,

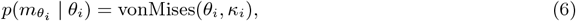

where the concentration parameter *κ*_*i*_ is converted from sensory noise 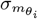 by using mean resultant length, 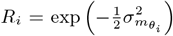, as defined by Fisher [88]. In the implementation, we use the approximation proposed by Best and Fisher [89], which has been shown to provide accurate estimates for practical applications:

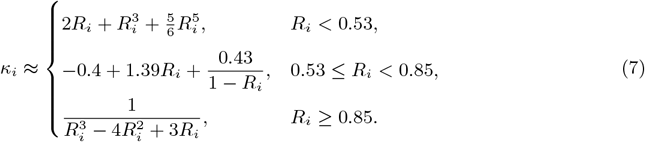

In contrast to the localization model, the likelihood of orientation is the same as the symmetric measurement distribution because sensory noise is conditioned on the measurement of the stimulus location, which is independent of the stimulus orientation. Assuming a uniform prior distribution, the posterior is proportional to the likelihood and symmetric. Consequently, the MAP and BLS estimators are the same as the orientation measurement, 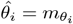.

Finally, we assume that the combined motor and adjustment noise is negligible compared with the sensory noise, and the orientation-estimation response is the same as the orientation estimate, 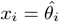.

The orientation-estimation model has two free parameters, Θ_*x*_ = *{a, b}*.

### 4.5 Confidence models

We denote the true stimulus value on a trial *i* by *u*_*i*_. For the localization task, *u*_*i*_ = *s*_*i*_ is the true stimulus location; for the orientation-estimation task, *u*_*i*_ = *θ*_*i*_ is the true orientation. In the second-order task of each trial *i*, the observer sets a confidence range *r*_*i*_ centered on their first-order estimate *û*_*i*_. The range width determines the potential reward according to the experimenter’s payoff function,

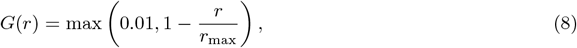

where *r* is the reproduced confidence range, and *r*_max_ is a fixed constant corresponding to the smallest potential reward of 0.01. In the localization task, *r*_max_ = 16.58 dva, one fourth of the screen width. In the orientation-estimation task, 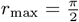 radians.

For Bayesian confidence models, we assume the same posterior distribution is used for the first-order response and the second-order computation of confidence [21, 22, 24].

#### 4.5.1 Ideal-observer confidence model

The ideal-observer model assumes the observer chooses the confidence range *r*_*i*_ to maximize expected subjective gain, using the posterior and a subjective-gain function,

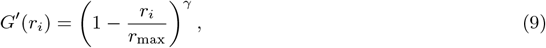

where the exponent *γ* is a risk-tolerance factor. *γ >* 1 indicates a convex subjective-gain function and a preference for smaller confidence range *r*.

The expected subjective gain of the trial *i* is:

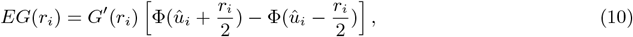

where Φ is the CDF of the posterior, *p*(*u*_*i*_ | *m*_*i*_), and *û*_*i*_ is the observer’s point estimate (MAP or BLS, depending on the first-order model).

The optimal confidence range is:

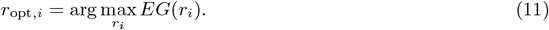

Observed confidence responses *c*_*i*_ are modeled as independent and identically distributed random variables from a log-normal distribution with mean *r*_opt,*i*_ and standard deviation *σ*_*c*_ (confidence noise and setting noise), guaranteeing that *c*_*i*_ will be positive.

The ideal-observer model has two free parameters Θ_*c*_ = *{γ, σ*_*c*_*}*.

#### 4.5.2 Alternative confidence models

Alternative models specify different computations of the confidence range *r*_*i*_ before noise is added, but otherwise share the same log-normal noise model for *c*_*i*_ (replacing the mean of the response distribution *r*_opt,*i*_ with *r*_*i*_). All models have two free parameters Θ_*c*_ = *{k, σ*_*c*_*}*. We start by introducing a heuristic model that does not use the posterior nor the gain function, followed by two Bayesian models without a subjective-gain function.

##### Scaled-measurement model

The scaled-measurement model assumes that the observer uses the distance that the location measurement *m*_*i*_ has from the center of the screen as a heuristic cue of confidence. Thus, the confidence range, *r*_*i*_, scales the magnitude of the location measurement *m*_*i*_ by a positive factor of *k*,

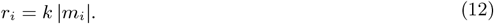

##### Scaled-posterior-width model

In the next two models, the observer applies a monotonic function to the posterior width in each trial, *σ*_post,*i*_, which represents the uncertainty of the location estimate, to obtain a confidence range. Note that any quantity that summarizes the precision of the posterior distribution can be a candidate for confidence range, such as entropy, interquartile range, and the probability density evaluated at the location estimate. These models should yield qualitatively similar predictions, and we focus on posterior width because it has the same unit as the confidence range *r*_*i*_ and thus is easier to interpret.

In the scaled-posterior-width model, the confidence range, *r*_*i*_, is proportional to the SD of the posterior distribution,

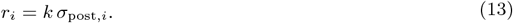

##### Scaled-log-posterior-width model

In this model, the confidence range, *r*_*i*_, is a scaled logarithmic function of the posterior width that ensures a positive prediction of confidence range,

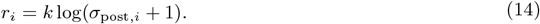

#### 4.5.3 Simplifications

We made two simplifying assumptions in our modeling. First, in the orientation-estimation model, we assumed that orientation noise depended on the true stimulus location. Strictly speaking, observers have access only to a noisy measurement, not the true stimulus location. We adopted this simplification to avoid introducing additional free parameters for spatial localization. Moreover, because the expected value of the measurement equals the stimulus, this approximation is reasonable for our purposes. Second, we did not include an orientation prior that could account for biases in orientation estimates. Although the observer might have less sensory noise or higher confidence for cardinal orientations, such effects should average out across trials at each eccentricity.

### 4.6 Model fitting and comparison

#### 4.6.1 Model likelihoods

We fit all models by maximizing likelihood of the observed data in two steps. First, we fit each observer’s first-order model to their localization or orientation-estimation responses. After obtaining the maximum-likelihood parameters governing first-order perception, we held these parameters fixed and then fit each confidence model to the same observer’s confidence data.

We denote the observed response on trial *i* by

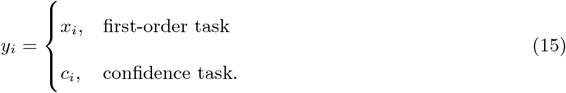

Let 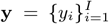 denote the full dataset for an observer. Assuming independent trials, the log likelihood of a model *M* and a parameter set Θ given the observed data is the sum of individual log likelihood across *I* trials,

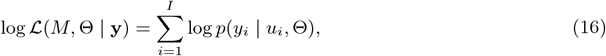

where *u*_*i*_ denotes the true stimulus value (*s*_*i*_ or *θ*_*i*_) on trial *i*.

For several models we considered, the likelihood *p*(*y*_*i*_ | *u*_*i*_) does not have a closed-form expression. In the confidence models, this arises because confidence is determined by an internal maximization of expected gain; in the localization models, the asymmetry of the sensory likelihood leads to analytically intractable posteriors. Therefore, we approximated the likelihood via numerical marginalization over the unobservable sensory measurement.

For each stimulus location *s*_*i*_, we discretized the internal measurement space into 2,000 linearly spaced samples over [-40, 40] dva. For each hypothetical measurement *m*_*i,j*_, the model predicts a first-order response *x*_*i,j*_ or a confidence range *c*_*i,j*_. Using the unified notation of responses *y*_*i,j*_, the likelihood on a given trial is approximated as

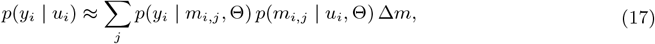

where Δ*m* is the grid spacing.

#### 4.6.2 Parameter estimation and model comparison

We used the BADS toolbox [90] in `MATLAB` to optimize the set of parameters for the models. We repeated each search 100 times with a different and random starting point to address the possibility of reporting a local minimum, and chose the parameter estimates with the maximum likelihood across the repeated searches.

For model comparison, we first computed AIC of each model by 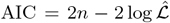, where *n* is the number of free parameters of the corresponding model [69]. We then calculated ΔAIC by subtracting the lowest AIC across models for each participant. We considered a participant best captured by a model if its ΔAIC *<* 2. Occasionally, one observer can have multiple best-fitting models that satisfy this criterion. Second, we evaluated predictive performance using five-fold cross-validation. For each participant, we partitioned the trials into five folds randomly, fit the model to four folds, and computed the negative log likelihood on the held-out fold. The cross-validated score was obtained by averaging across folds.

## 5 Acknowledgments

All data and code are available via the Open Science Framework (https://osf.io/5cgbt) after acceptance. This research was supported by NIH grant EY08266, NIH Training Program in Computational Neuroscience (TPCN) grant T90DA059110, and the NYUAD Center for Brain and Health, funded by Tamkeen under NYU Abu Dhabi Research Institute grant CG012. This work was also supported in part through the NYU IT High Performance Computing resources, services, and staff expertise.

