## Supplementary Information for "Visual confidence accurately tracks increasing internal noise with eccentricity in peripheral vision"

Luhe Li[1] and Michael S. Landy[1,2]

[1] *Department of Psychology, New York University*

[2] *Center for Neural Science, New York University*

### Appendix 1 Comparing spatial localization models

In this section, we describe four models of spatial localization, followed by the results of model comparison.

#### Appendix 1.1 Methods

We considered localization models that were a factorial combination of two modeling choices. The first factor concerns how spatial bias is represented. Bias can be modeled either by a central prior, such as a zero-mean Gaussian distribution, $p(s_i) = \mathcal{N}(0, \sigma_s^2)$, or by a linear remapping from physical space to internal space, $s_i' = \beta\, s_i$. The Gaussian prior captures only a bias toward the center, whereas linear remapping can account for biases toward either the center or the periphery by setting $\beta < 1$ or $\beta > 1$, respectively.

The second factor concerns the estimator used to read out the posterior distribution: either its mean (the Bayesian least-squares estimator, BLS) or mode (the maximum-a-posteriori estimator, MAP). Usually these two estimators are the same when the posterior is a symmetric Gaussian, but when sensory noise grows with stimulus magnitude, the likelihood and posterior are asymmetric and the two estimators can differ.

We fit each observer's localization responses by maximizing likelihood using the same procedures described in the Methods section. We evaluated model performance using the Akaike Information Criterion (AIC). We calculated $\Delta$AIC by subtracting the lowest AIC across models for each participant. We considered a participant best captured by a model if its $\Delta\text{AIC} < 2$. Occasionally, one observer can have multiple best-fitting models that satisfy this criterion.

#### Appendix 1.2 Results

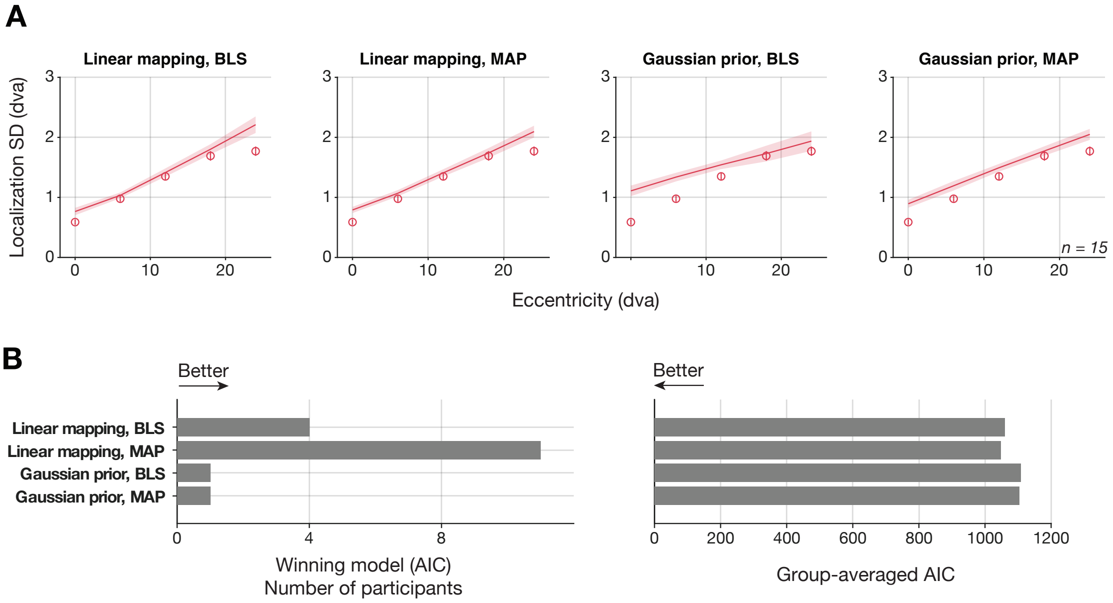


Figure S1: Model prediction and comparison. (A) Model prediction of localization standard deviation (SD) as a function of eccentricity. Group-averaged data (dots, $n = 15$) and model prediction (lines and shades) of each localization model. The linear-mapping models (left two panels) accurately capture localization variability at the fovea, whereas the Gaussian-prior models (right two panels) consistently overestimate the variability at the fovea. Error bar and shades: $\pm 1$ SEM across observers. (B) Quantitative comparison using AIC. The linear-mapping model provided the best fit for the majority of participants (i.e., $\Delta \text{AIC} < 2$) and has lower group-averaged AIC values.

The model comparison revealed a decisive preference for the linear-mapping model over the Gaussian-prior model across the participants. Qualitatively, model predictions showed that Gaussian-prior models consistently overestimated localization variability at the fovea, whereas linear-mapping models provided a better fit across eccentricities (Fig. S1A). Quantitatively, the linear-mapping model was the winning model for 14 out of 15 participants at the individual level (Fig. S1B). This result was corroborated by group-averaged AIC values, which indicated that linear-mapping models consistently outperformed Gaussian-prior models.

Based on these results, we selected the linear-mapping model as the foundational localization model. We adopted an individual-best-model approach to account for idiosyncratic differences between observers, as some were best fit by the BLS estimator and others by the MAP estimator. Therefore, for all subsequent confidence modeling, we used the specific estimator within the linear-mapping model that minimized the AIC for each individual participant.

### Appendix 2 Individual-level model predictions of the spatial localization and confidence models

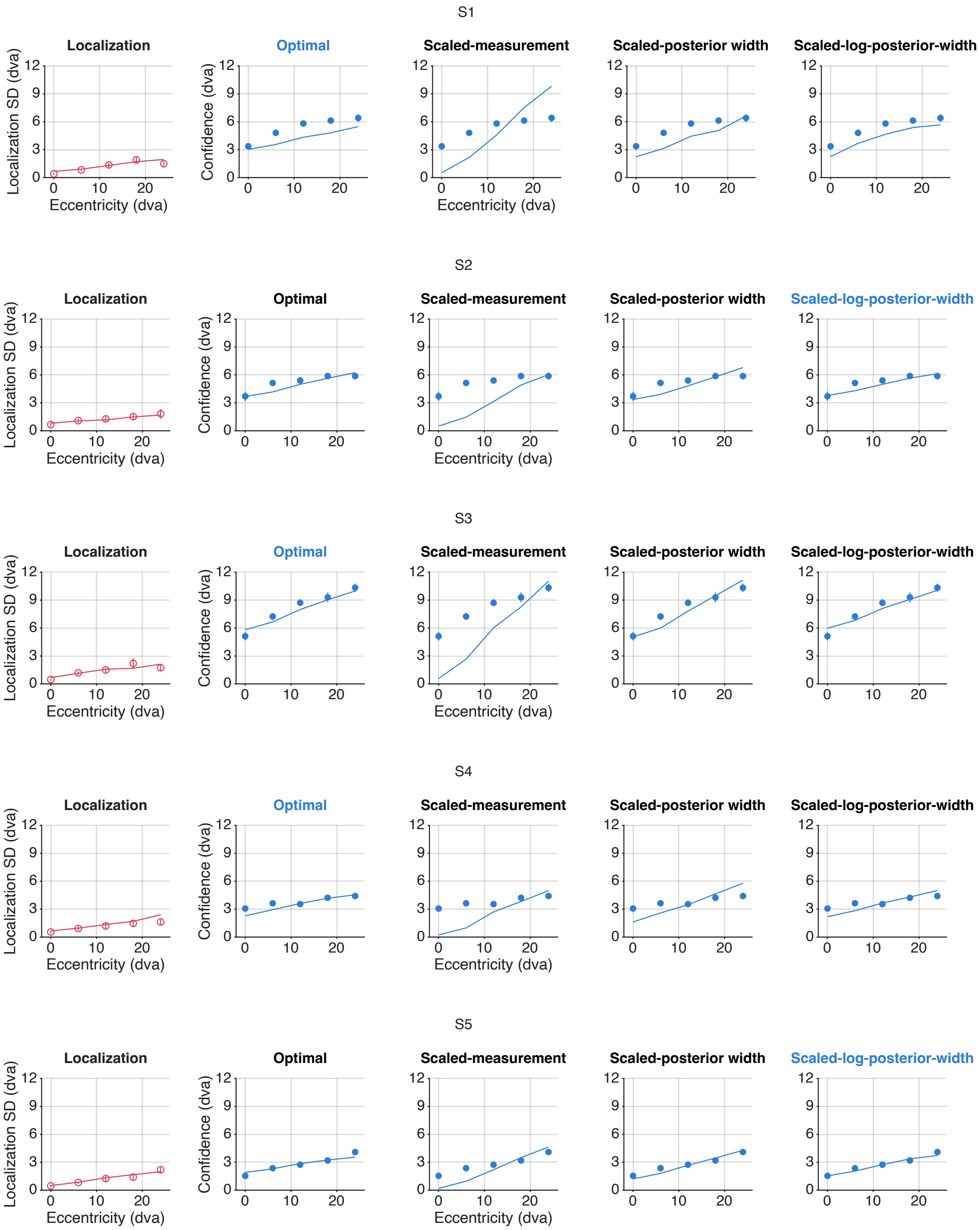


Figure S2: Individual-level data (dots) and model predictions (lines) of the spatial localization and confidence models. Error bars are 68% confidence interval of 10000 bootstrapped trials. Blue error bars can be invisible when they are smaller than the data points. The localization-model prediction (red) is the same across confidence models (blue). The title of the best confidence model for each individual participant based on AIC comparison is highlighted in blue.

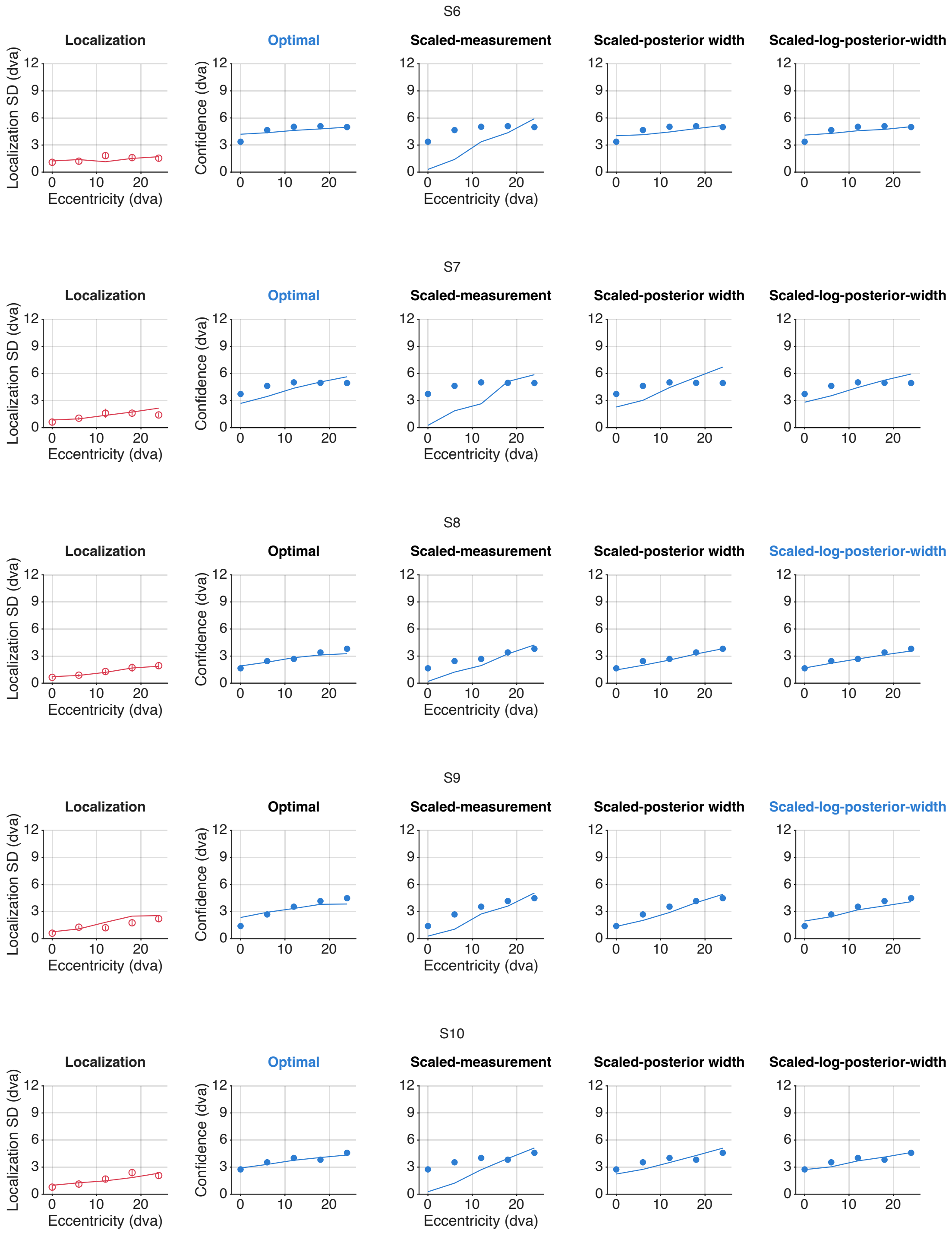


Figure S2 (continued).

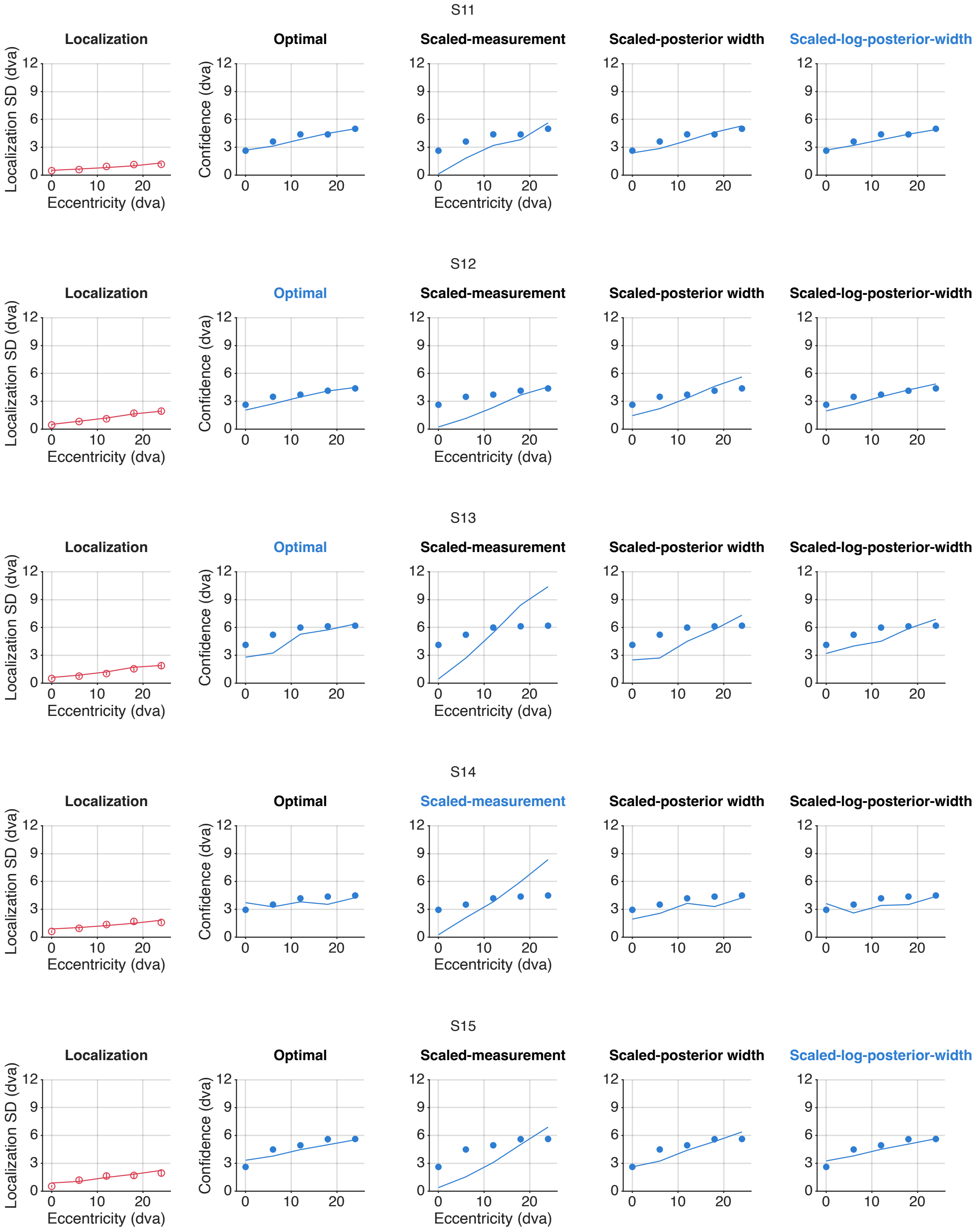


Figure S2 (continued).

### Appendix 3 Individual-level parameter estimates of winning models (Experiment 1: spatial localization task)

Table S1: Parameter estimates of the ideal-observer spatial localization model.

| | a (Scaling factor) | b (Intercept) | $\beta$ (Spatial-bias scaler) |
|---|---|---|---|
| S1 | 0.063 | 0.612 | 0.947 |
| S2 | 0.041 | 0.749 | 0.967 |
| S3 | 0.064 | 0.858 | 0.876 |
| S4 | 0.077 | 0.510 | 0.946 |
| S5 | 0.073 | 0.482 | 0.931 |
| S6 | 0.021 | 1.294 | 0.928 |
| S7 | 0.054 | 0.556 | 1.115 |
| S8 | 0.063 | 0.583 | 0.942 |
| S9 | 0.107 | 0.682 | 0.901 |
| S10 | 0.063 | 0.908 | 0.946 |
| S11 | 0.032 | 0.494 | 0.953 |
| S12 | 0.074 | 0.442 | 0.940 |
| S13 | 0.073 | 0.462 | 0.937 |
| S14 | 0.040 | 0.835 | 0.968 |
| S15 | 0.062 | 0.802 | 0.914 |
| Lower bound | 0.010 | 0.100 | 0.200 |
| Upper bound | 0.500 | 5.000 | 1.800 |
| Group mean | 0.061 | 0.685 | 0.947 |
| Group SEM | 0.001 | 0.015 | 0.003 |

Table S2: Parameter estimates of the ideal-observer confidence model.

| | $\beta$ (Exponent) | $\sigma_c$ (Confidence noise) |
|---|---|---|
| S1 | 1.034 | 2.351 |
| S2 | 0.468 | 0.269 |
| S3 | 0.093 | 0.191 |
| S4 | 1.771 | 0.229 |
| S5 | 2.798 | 0.321 |
| S6 | 1.111 | 0.226 |
| S7 | 0.954 | 0.254 |
| S8 | 3.164 | 0.279 |
| S9 | 2.732 | 0.381 |
| S10 | 2.112 | 0.211 |
| S11 | 0.446 | 0.274 |
| S12 | 1.727 | 0.265 |
| S13 | 0.627 | 2.002 |
| S14 | 1.820 | 1.985 |
| S15 | 1.106 | 0.321 |
| Lower bound | 0.010 | 0.010 |
| Upper bound | 5.000 | 4.000 |
| Group mean | 1.464 | 0.637 |
| Group SEM | 0.062 | 0.051 |

Table S3: Parameter estimates of the scaled-log-posterior-width confidence model.

| | $\beta$ (Exponent) | $\sigma_c$ (Confidence noise) |
|---|---|---|
| S1 | 2.567 | 2.355 |
| S2 | 3.097 | 0.268 |
| S3 | 4.248 | 0.193 |
| S4 | 2.059 | 0.270 |
| S5 | 1.614 | 0.305 |
| S6 | 2.441 | 0.228 |
| S7 | 2.681 | 0.263 |
| S8 | 1.588 | 0.270 |
| S9 | 1.437 | 0.365 |
| S10 | 1.861 | 0.222 |
| S11 | 3.081 | 0.273 |
| S12 | 2.099 | 0.298 |
| S13 | 2.891 | 2.004 |
| S14 | 2.029 | 1.985 |
| S15 | 2.453 | 0.319 |
| Lower bound | 0.010 | 0.010 |
| Upper bound | 5.000 | 4.000 |
| Group mean | 2.410 | 0.641 |
| Group SEM | 0.049 | 0.051 |

### Appendix 4 Comparison of confidence models built on an alternative model of orientation estimation

To assess whether the superior performance of the ideal-observer model was robust to assumptions about sensory noise, we tested an alternative orientation-estimation model in which noise increased as an exponential function of eccentricity. Under this formulation, Eq. 5 becomes $\sigma_{m_{\theta_i}} = e^{(a\,|s_i|+b)}$. This allows the scaled-posterior-width model to predict a nonlinear increase in confidence with eccentricity.

We fitted the orientation-estimation responses by maximum likelihood following the procedure in the Methods and fixed the resulting sensory-noise parameters for each observer. Finally, we re-evaluated the suite of confidence models described in the main text.

The model comparison yielded results consistent with the primary analysis of Experiment 2. Even with the assumption of exponentially growing sensory noise, the ideal-observer model provided a better fit to the confidence data. Quantitatively, the ideal-observer model was the preferred model for the majority of participants (i.e., $\Delta$AIC relative to the smallest AIC is less than 2, thus one observer can have multiple winning models) and achieved the lowest group-averaged AIC.

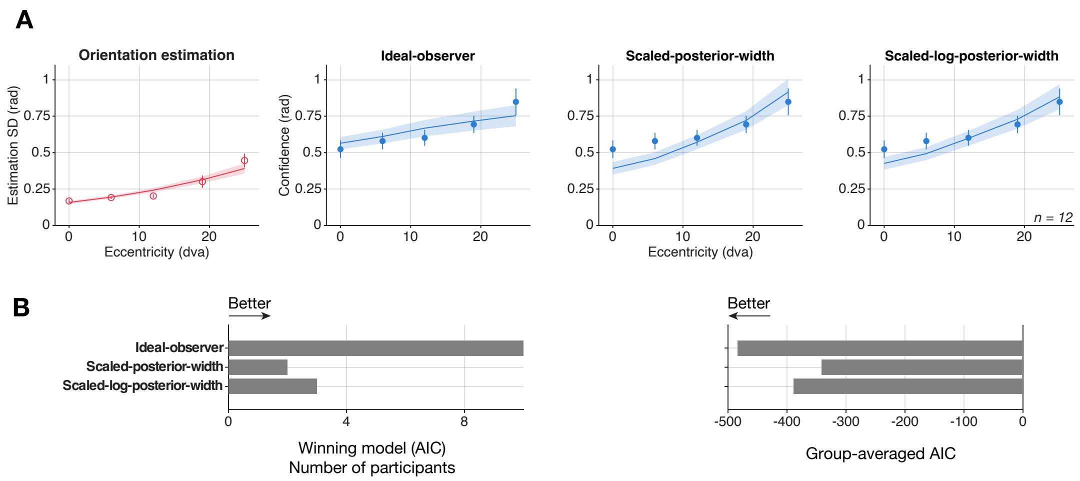


Figure S3: Model prediction and comparison. (A) Group-averaged data (dots, $n = 12$) and model prediction (lines and shades). The orientation-estimation model prediction (red) is fixed across all panels. With an exponentially-increasing sensory noise, the ideal-observer model (left-most blue panel) still better captures the data than the rest of the models. Error bar and shades: $\pm 1$ SEM across observers. (B) Quantitative comparison using AIC. The ideal-observer model captured the majority of the participants (i.e., $\Delta\text{AIC} < 2$) and had the lowest group-averaged AIC values.

### Appendix 5 Individual-level model predictions of the orientation-estimation and confidence models

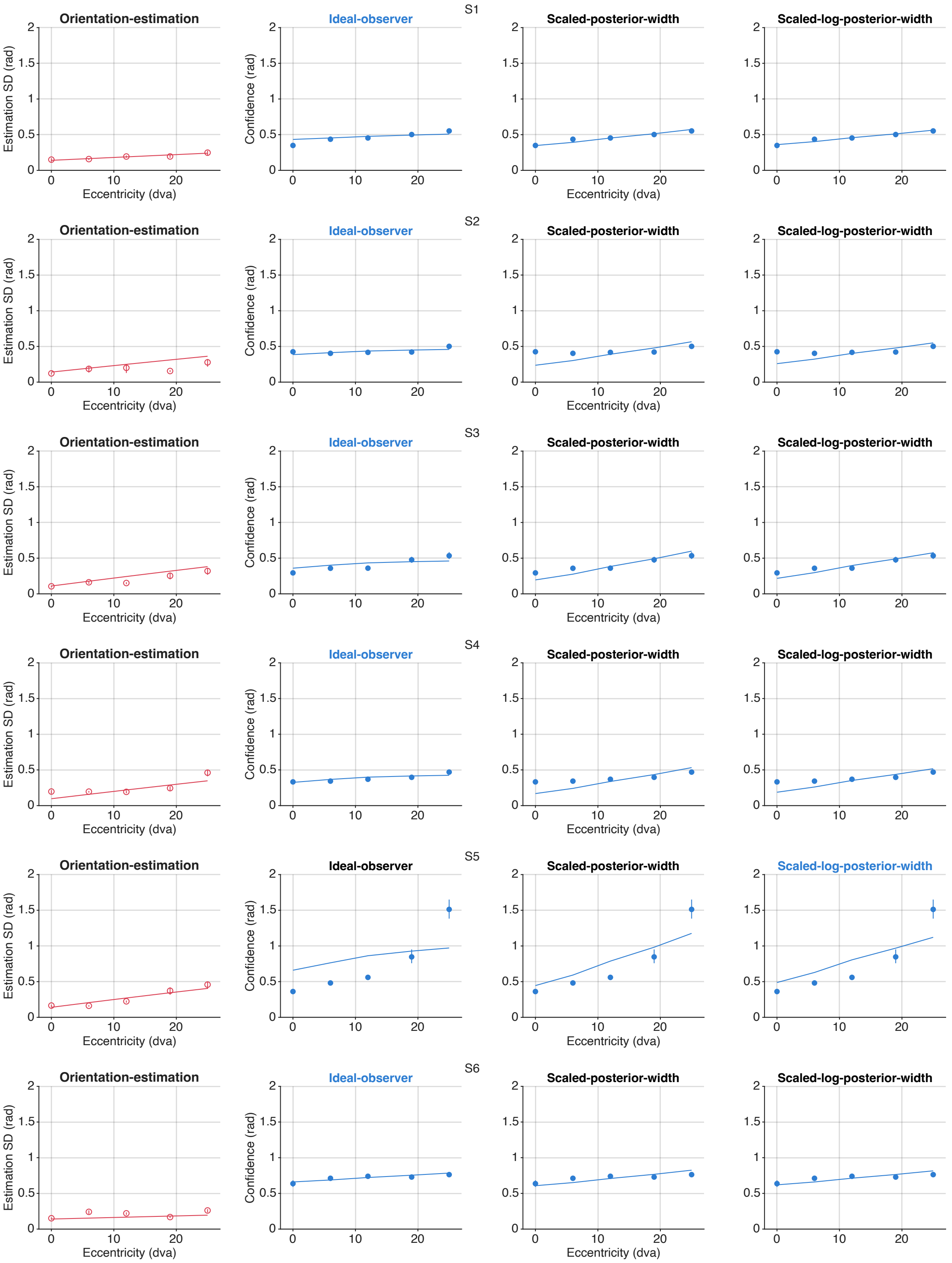


Figure S4: Individual-level data (dots) and model predictions (lines) of the orientation-estimation and confidence models. Error bars are 68% confidence interval of 10000 bootstrapped trials. Blue error bars can be invisible when they are smaller than the data points. The orientation-estimation-model prediction (red) is the same across confidence models (blue). The title of the best confidence model for each individual participant based on AIC comparison is highlighted in blue.

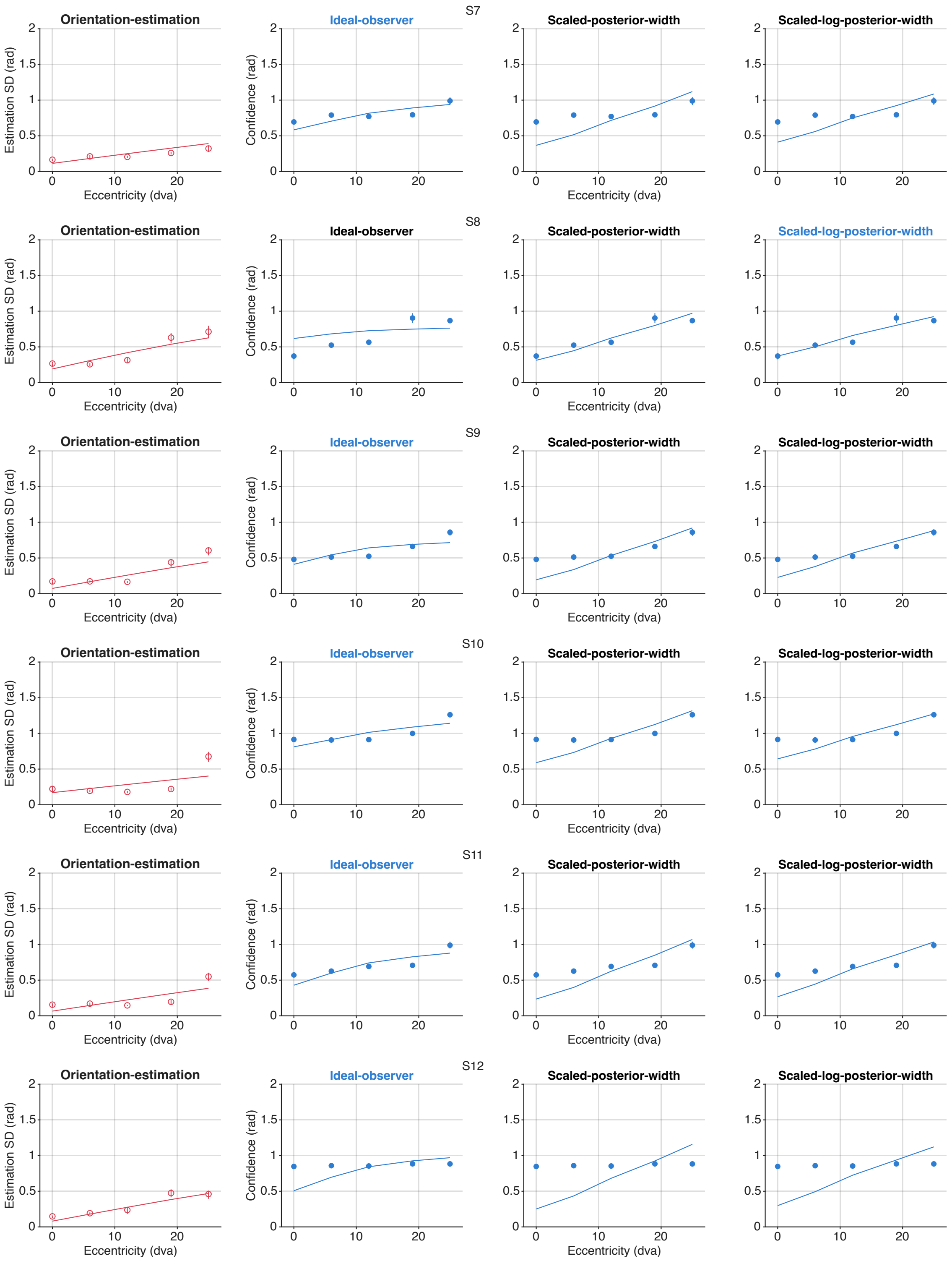


Figure S4 (Continued).

### Appendix 6 Individual-level parameter estimates of winning models (Experiment 2: orientation-estimation task)

Table S4: Parameter estimates of the ideal-observer orientation-estimation model.

| | a (Scaling factor) | b (Intercept) |
|---|---|---|
| S1 | 0.006 | 0.200 |
| S2 | 0.013 | 0.200 |
| S3 | 0.016 | 0.156 |
| S4 | 0.015 | 0.137 |
| S5 | 0.016 | 0.200 |
| S6 | 0.003 | 0.200 |
| S7 | 0.016 | 0.161 |
| S8 | 0.029 | 0.273 |
| S9 | 0.022 | 0.104 |
| S10 | 0.014 | 0.242 |
| S11 | 0.019 | 0.094 |
| S12 | 0.024 | 0.115 |
| Lower bound | 0.001 | 0.010 |
| Upper bound | 0.050 | 0.500 |
| Group mean | 0.016 | 0.173 |
| Group SEM | 0.001 | 0.005 |

Table S5: Parameter estimates of the ideal-observer confidence model.

| | $\beta$ (Exponent) | $\sigma_c$ (Confidence noise) |
|---|---|---|
| S1 | 4.341 | 0.065 |
| S2 | 5.564 | 0.071 |
| S3 | 5.559 | 0.106 |
| S4 | 6.104 | 0.057 |
| S5 | 1.731 | 0.332 |
| S6 | 1.477 | 0.124 |
| S7 | 1.824 | 0.138 |
| S8 | 2.950 | 0.141 |
| S9 | 3.051 | 0.139 |
| S10 | 1.218 | 0.118 |
| S11 | 2.058 | 0.121 |
| S12 | 1.845 | 0.102 |
| Lower bound | 0.001 | 0.0001 |
| Upper bound | 10.000 | 20.000 |
| Group mean | 3.143 | 0.126 |
| Group SEM | 0.149 | 0.006 |
